## Supplementary methods and figures for "Substitution load revisited: a high proportion of deaths can be selective"

### SUPPLEMENTARY INFORMATION

#### Supplementary Methods

##### Environmental conditions

We re-analyze the data of Exposito-Alonso et al. (2019), who used a 2 × 2 × 2 design for environmental conditions, with the three treatments being climate, water availability, and adult density (see github for raw data and analysis code). We treat these as eight separate populations. For climate, plants were grown in outdoor field stations in either Tübingen, Germany (near the center of the species range of *A. thaliana*) or in Madrid, Spain (at the southern edge of the range). Plants were all artificially watered. The high-water treatment matched soil moisture levels near the station in Germany, and the low-water treatment matched soil moisture levels near the station in Spain. To generate high adult density, thirty seeds of the same genotype were planted per pot. For low density, several seeds (~10) were planted per pot, enough to ensure that at least one seed would germinate, but few enough that the seeds were unlikely to inhibit each other pre-germination. To avoid any competition between adult plants in the low density treatment, only one seedling, chosen at random, was retained after germination, and the rest were plugged out and discarded. We refer to each treatment with a three-letter abbreviation: M or T for Madrid or Tübingen, L or H for low or high water, and I or P for a single individual plant or a population of thirty plants per pot. For example, the treatment with thirty seeds per pot grown in Madrid with high water is abbreviated as MHP.

Within each of the eight populations, seeds from 517 fully homozygous plant genotypes (taken from a parental generation grown under controlled conditions to control for parental effects) were grown in pots that included only plants of that genotype. The number of replicate pots per genotype per population was occasionally as few as one due to experimental losses, but mostly ranged between five to seven replicates. Our interest is in differences among genotypes, not among replicates. We therefore calculate key quantities of interest for each genotype-environment combination by averaging across replicates.

##### Selective deaths

The experiment can be mapped reasonably easily onto theoretical treatments. In each environmental treatment, a starting population of seeds grows into adult plants, experiencing both selective deaths and non-selective deaths as they proceed from seeds to seedlings to adults. Plants which survive to become adults then produce $k$ seeds on average, some of which would normally constitute the next generation, although the experiment concludes at the end of season. The experiment does not capture the life history stage of seed dispersal to fertile ground, to complete the life cycle that began with seeds planted in a pot.

Juvenile deaths must be treated differently for the low- and high-density treatments. In the low-density treatment, where exactly one seedling is retained after germination, we do not have access to data on selective seed deaths, and so consider only seedling selective deaths. In the high-density treatment, any of the thirty seeds that fail to survive to the end of the experiment are counted as deaths, whether due to seed death before germination or to subsequent seedling death; our selective death calculations do not differentiate between these two life history transitions. This means that across the two life history transitions at which juvenile plants can die (as planted seeds before germination and as seedlings), only one set of juvenile deaths is recorded in each density treatment, but they are not comparable. They are combined seed and seedling deaths in the high-density case, and seedling deaths alone in the low-density case. Histograms are shown in Supplementary Figures 1-2.

In each treatment, we score the observed juvenile death rate of the highest performing genotype as the baseline extrinsic mortality for all genotypes (i.e. as non-selective deaths). Conceptually (ignoring a correction for extreme value bias, treated below), for each life history transition in each environmental condition we have:

$$\text{Selective deaths in the population } = \sum_i n_i(d_i - d_{best})$$

where $n_i$ is the starting population of genotype $i$ at that life history transition, $d_i$ is the genotype's average death rate during that life history transition, and $d_{best}$ is the average death rate of the genotype with the lowest death rate for that life history transition in that environment.

#### Test for genetic variance in fecundity

Selective "deaths" can also be defined for unrealized fecundity. We did not analyze this here, because of lack of evidence for significant genetic differences in fecundity. In support of this, we performed an ANOVA test on fecundity in each environmental condition. We only have information on fecundity as an aggregate per replicate pot (rather than per individual plant in the high-density condition), so we compare among-genotype variance to among-replicate variance. Note, however, that we remove replicates that had no adults surviving to reproductive maturity, as well as genotypes with only a single replicate pot with surviving adults (and therefore no way of estimating among-replicate variance). All surviving adults produced at least some seeds. We Box-Cox transformed the data for each pot with surviving adults in each environmental condition (see Supplementary Figure 3 for post-transform histograms) before performing the ANOVA.

#### Proportion of juvenile deaths selective

Because *A. thaliana* is an annual plant, all juveniles will die by the end of the season, whether as selective deaths during the experiment, non-selective deaths during the experiment, or non-selective deaths after the end of the experiment. From this we obtain, for each environmental condition:

$$\text{Fraction of juvenile deaths selective} = \frac{Selective\ deaths\ of\ juveniles}{Starting\ population}$$

In the high-density populations, the starting population is 30 seeds per pot. In the low-density populations, the starting population is 1 seedling per pot.

#### Pairwise genotype comparisons

For every possible pair of genotypes, we repeat the analysis above to estimate selective deaths and the proportion of deaths which are selective, using the better genotype of the pair as the 'best' genotype in the calculation of selective deaths. With only two genotypes, we do not adjust for extreme value bias. Using whole-genome information, we calculated the total number of SNP differences between each pair (Hamming distance, number of allele differences out of 1,353,386 biallelic SNPs) using PLINK v1.9.

#### Correction for extreme value bias

The estimated best genotype is subject to extreme value bias, leading to overestimation of the number of selective deaths. I.e., the best genotype observed is likely not only to be a superior genotype, but also to have outperformed its own expected death rate by chance. The more uncertainty in estimated genotypic survival rates, relative to true genetic variance, the worse the extreme value bias problem. Here we attempt a rough estimate of the magnitude of extreme value bias using the observed noise among replicates and among genotypes. We then subtract a conservative estimate of bias from our best observed genotype, and redo our calculations of selective deaths.

We perform 10,000 simulations per environmental treatment. In each simulation, we assume that the observed 517 genotypic death rates are the 'true' values for each genotype (thus slightly overestimating genetic variance within the population). We then resample the number of surviving seeds 'observed' for each replicate of that genotype using a binomial distribution and calculate the resulting observed death rate.

In each simulated dataset, we took the best observed genotype and recorded the difference between its observed value and its actual genotypic value, then averaged these differences across the 10,000 simulations to obtain estimated bias. We then adjusted our estimate of the number of selective deaths to incorporate this estimated bias:

$$\text{Total adjusted selective deaths} = \sum_{i=1}^{517} n_i(d_i - (d_{best} + bias)).$$

Adjustments to selective deaths are shown in Supplementary Table 1. Across all environmental treatments, adjusting for extreme value bias leads to negligible change in the estimate of the proportion of deaths which are selective.

In six out of eight environmental conditions, at least one genotype had all plants survive until the end of the experiment. This genotype then has an observed death rate of zero and an observed between-replicate variance of zero, and thus our approach will not find any bias. Determining bias in this case would require attempting to fit and then resample from a true distribution of genotypic values that includes death rates near but not equal to zero. We did not pursue this avenue, because resampling genotypes would add a new source of variance, and we need only correct for the extreme value bias pertaining to the genotypes actually studied. Given the small degree of bias observed in the two environmental conditions in which no genotype experienced perfect survival, we consider our approach sufficient to conclude that extreme value bias has little quantitative effect on our results.

#### Reproductive excess calculation

The experiment analyzed here has a fixed life cycle of four life history transitions (adults producing seeds, seeds successfully dispersing to suitable habitat, seeds surviving to be seedlings, seedlings surviving to be adults). Matching this, we define reproductive excess ($RE$) four different ways, as illustrated in Figure 3. Given the presence of many poorly adapted genotypes in the experiment, we perform each calculation with respect to the best genotype (denoted by the prime symbol $'$), yielding:

$$\text{RE(seed survival)} = k'_{seed\_survival}N_{seeds} - \frac{N_{seeds}}{k'_{seedling\_survival}k'_{fecundity}k'_{dispersal}}$$

$$\text{RE(seedling survival)} = k'_{seedling\_survival}N_{seedlings} - \frac{N_{seedlings}}{k'_{fecundity}k'_{dispersal}k'_{seed\_survival}}$$

$$\text{RE(fecundity)} = k'_{fecundity}N_{adults} - \frac{N_{adults}}{k'_{dispersal}k'_{seed\_survival}k'_{seedling\_survival}}$$

$$\text{RE(dispersal)} = k'_{dispersal}N_{seeds\_produced} - \frac{N_{seeds\_produced}}{k'_{seed\_survival}k'_{seedling\_survival}k'_{fecundity}}$$

In the high density environmental conditions, we use a single survival transition to cover both seeds and seedlings. Under low density conditions, where one seedling was chosen at random from the product of 10 planted seeds, we use $k_{seed\_survival} = 0.1$ for all genotypes. We treat all genotypes as having the same $k_{fecundity}$ (due to lack of evidence for genetic variation — see Experimental Results below). Our experiment provides no information about values of $k_{dispersal}$, so we consider a range from 0.01 to 0.1, equal across genotypes — this uncertainty produced by this range is represented in Supplementary Table 2, and by the vertical lines in Figure 4. Because we lack data on the number of seeds after dispersal, we do not calculate a reproductive excess for this transition. This means that we calculate only two reproductive excesses for each environmental condition, one for survival and one for fecundity, although the calculations are different between low- and high-density conditions. These reproductive excesses are given by:

$$\text{RE(fecundity, low density)} = k_{fecundity}N_{adults} - \frac{N_{adults}}{0.1k_{dispersal}k'_{seedling_{survival}}} \quad (1)$$

$$\text{RE(survival, low density)} = k'_{seedling\_survival}N_{seedlings} - \frac{N_{seedlings}}{0.1k_{dispersal}k_{fecundity}} \quad (2)$$

$$\text{RE(fecundity, high density)} = k_{fecundity}N_{adults} - \frac{N_{adults}}{k_{dispersal}k'_{survival}} \quad (3)$$

$$\text{RE(survival, high density)} = k'_{survival}N_{seeds} - \frac{N_{seeds}}{k_{dispersal}k_{fecundity}} \quad (4)$$

### Supplementary Results

#### Selection on fecundity

Our empirical findings are restricted to selective deaths relating to juvenile survival. We did not analyze selective 'deaths' attributable to differences in fecundity, because an ANOVA on fecundity within and among genotypes showed no statistical support for any difference in mean fecundity among genotypes in six of the eight experimental conditions. Even in the two conditions with statistical significance, among-genotypic variance was three times smaller than among-replicate variance. A more sensitive MCMCglmm model with Poisson errors and controlling for replicate found non-zero heritability in 6/8 environments, but still below 10%, which we consider low enough to neglect.

#### Dependence on environment

We expected *a priori* that harsher environmental conditions would have higher extrinsic mortality. However, this wasn't the case. We can estimate extrinsic mortality as the death rate of the best genotype, after correcting for extreme value bias (see columns 3 and 4 of Supplementary Table 1). Most environmental conditions saw a highest-fitness genotype with perfect survival, or close to it. We saw the most extrinsic mortality in the THP environmental condition, which is not one of the harsher conditions. Nei (1971) and Felsenstein (1971) implicitly assume high extrinsic mortality via their choice of value for reproductive excess; we explicitly account for extrinsic mortality during mortality transitions. Extrinsic mortality might of course be much higher in natural conditions, lowering the proportion of deaths below the high values observed here.

#### Dependence on genotypes

An obvious concern with our setup is that with genotypes representing Europe-wide diversity of *A. thaliana*, exaggerated differences between the best-adapted and worst-adapted genotypes would inflate estimates of the proportion of deaths that were selective. If this were the case, then we expect that competition between more similar genotypes should lead to a smaller estimate for this proportion. We tested this prediction by repeating our analysis on every pair of genotypes, as though they were the only two genotypes in the experiment, and looking for a correlation between genetic distance and proportion of deaths selective. Note that we use genetic distance, much of it presumably neutral, as a proxy for genetic differences related to adaptation. Although some statistically significant correlations were observed in some environmental conditions, the direction of correlation was evenly split between negative and positive (as seen in Supplementary Table 3), and the highest $R^2$ value observed in any environmental condition was $0.096^2 = 0.0092$, for seedling deaths in the MLI environment, which we deem biologically insignificant. This is reassuring with respect to the artificially high genetic diversity in our experiment. However, two similar genotypes in our experiment represent more genetic distance than might be present within a typical natural population, and even closely related genotypes might differ in important fitness-associated traits.

### Supplementary Tables

| Environmental condition | Among-genotype variance | Observed maximum survival rate | Estimated bias | Unadjusted proportion of deaths selective | Adjusted proportion of deaths selective |
|---|---|---|---|---|---|
| MLI | 0.0618 | 1 | 0 | 0.664 | 0.664 |
| MHI | 0.0144 | 1 | 0 | 0.0847 | 0.0847 |
| TLI | 0.0337 | 1 | 0 | 0.564 | 0.564 |
| THI | 0.0139 | 1 | 0 | 0.109 | 0.109 |

| | | | | | |
|---|---|---|---|---|---|
| MLP | 0.0113 | 1 | 0 | 0.96 | 0.96 |
| MHP | 0.0433 | 1 | 0 | 0.564 | 0.564 |
| TLP | 0.023 | 0.947 | 0.00418 | 0.633 | 0.629 |
| THP | 0.0195 | 0.871 | 0.0195 | 0.391 | 0.372 |

Supplementary Table 1: Adjusting for extreme value bias has little effect on the estimated proportion of selective deaths.

| | **Proportion of seedling deaths selective** | **Proportion of combined seed and seedling deaths selective** | **Excess seedlings per pot after seedling survival transition** | **Excess seedlings per pot after seed and seedling survival transition** | **Excess seeds produced per pot after fecundity transition** |
|---|---|---|---|---|---|
| **MLI** | 0.664 | | 0.742-0.974 | | 967-1,269 |
| **MHI** | 0.0847 | | 0.945-0.995 | | 15,856-16,680 |
| **TLI** | 0.564 | | 0.794-0.979 | | 1,683-2,075 |
| **THI** | 0.109 | | 0.908-0.991 | | 8,848-9,651 |
| **MLP** | | 0.960 | | 20.51-29.05 | 257-364 |
| **MHP** | | 0.564 | | 28.16-29.82 | 20,029-21,207 |
| **TLP** | | 0.633 | | 13.71-22.97 | 833-1,396 |
| **THP** | | 0.391 | | 21.52-24.47 | 10,374-11,796 |

Supplementary Table 2**.** The proportion of deaths that are selective is above Haldane's 10% estimate for most environmental conditions. Reproductive excess after the survival and fecundity life history transitions are high. Note that reproductive excess per pot for low-density conditions cannot exceed one, and excess per pot for high-density conditions cannot exceed 30. For reproductive excess, lower bounds are calculated using $k_{dispersal} = 0.01$ and higher bounds are calculated using $k_{dispersal} = 0.1$. Reproductive excess values for fecundity are rounded to the nearest integer; approximate methods were used to estimate seed number.

| Life history stage | Seedling deaths | | Combined seed and seedling deaths | |
|---|---|---|---|---|
| Environmental condition | Spearman's rho | p-value | Spearman's rho | p-value |
| MLI | 0.096 | 2.2e-16 | | |
| MHI | 0.0004 | 0.88 | | |
| TLI | -0.0024 | 0.39 | | |
| THI | -0.0056 | 0.042 | | |
| MLP | | | 0.055 | 2.2e-16 |
| MHP | | | 0.011 | 4.9e-5 |
| TLP | | | -0.0081 | 0.0031 |
| THP | | | -0.0085 | 0.002 |

Supplementary Table 3. Genetic distance between a pair of genotypes does not consistently correlate with proportion of deaths selective. Visualization of each of the eight relationships is available in Supplementary Figures 4-5.

#### Supplementary Figures

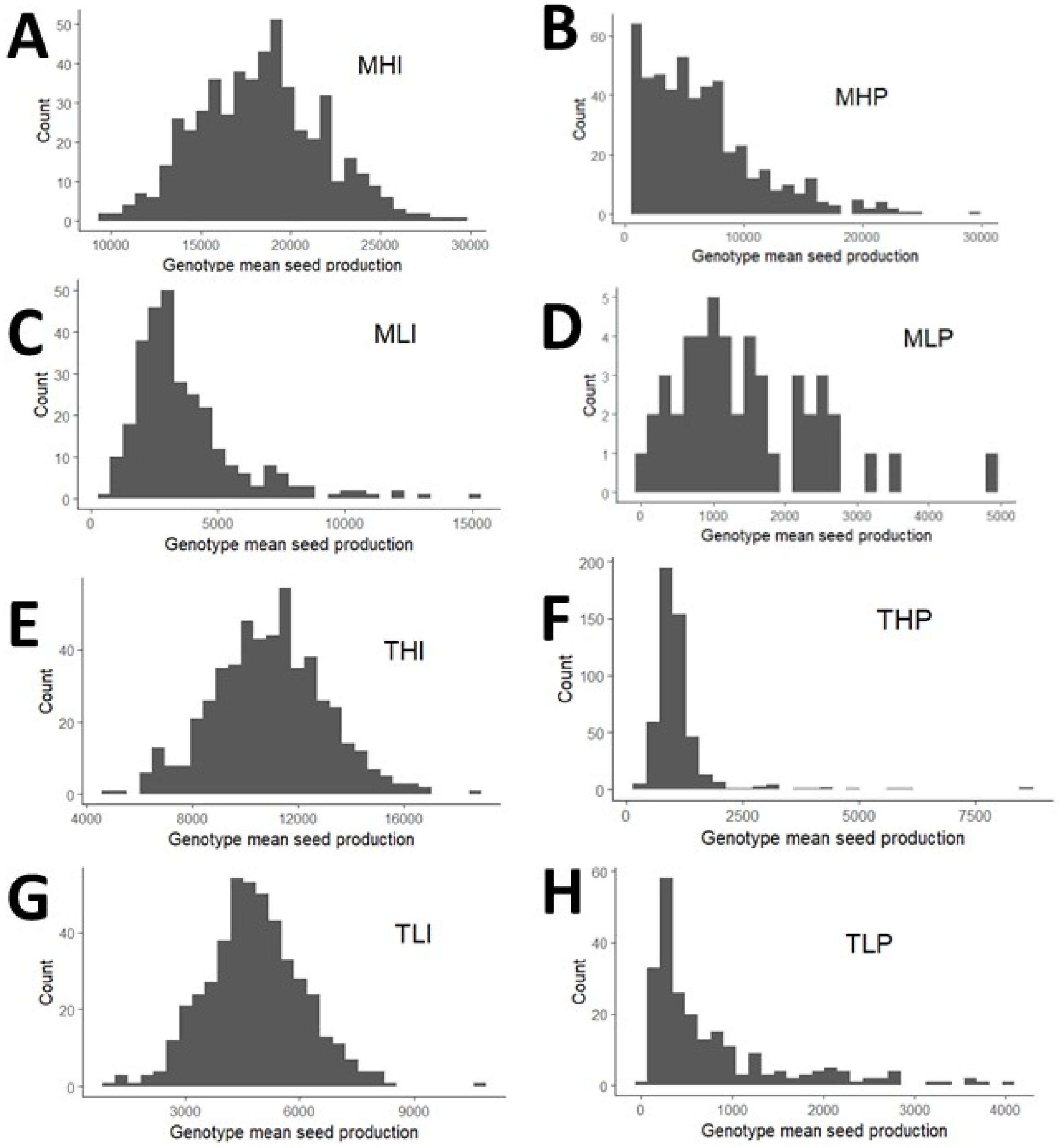


Supplementary Figure 1**.** Histograms of genotype mean seed production for every genotype with at least one surviving adult in each of the eight environmental conditions.

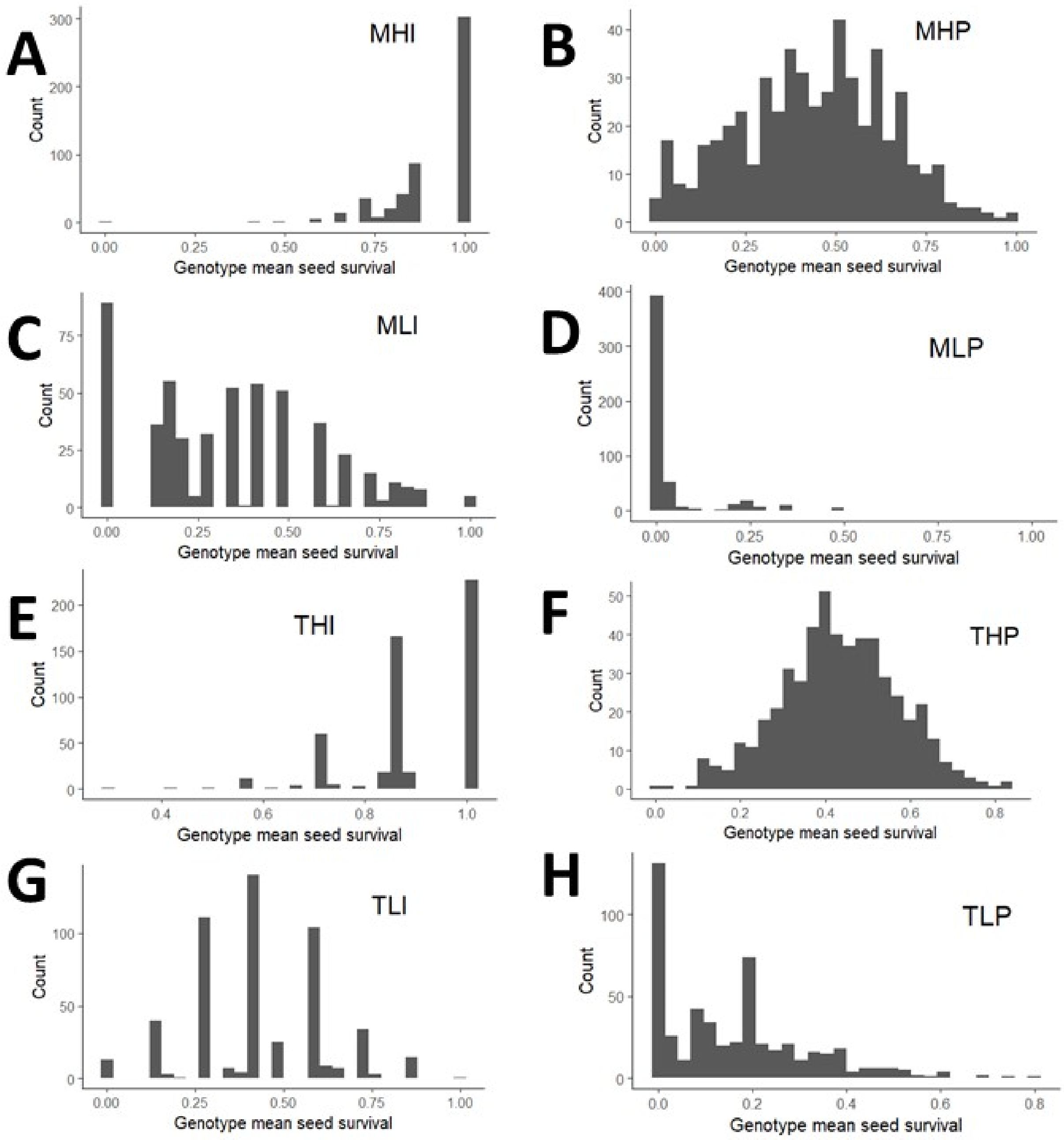


Supplementary Figure 2. Histograms of genotype mean seed survival for every genotype in each of the eight environmental conditions.

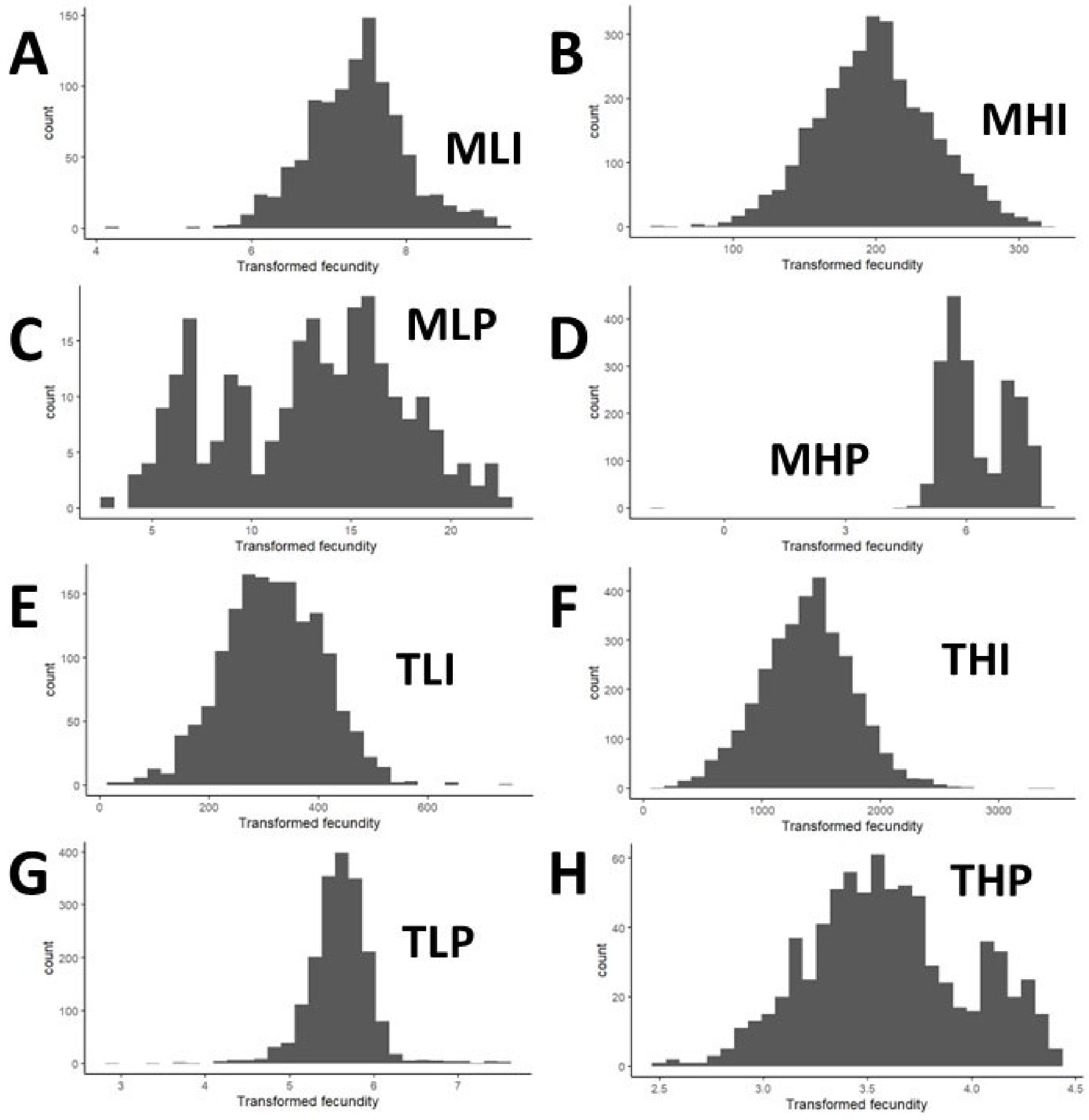


Supplementary Figure 3**.** Histograms of Box-Cox transformed fecundity values for every pot with surviving adults in each of the eight environmental conditions. Lambda values for the Box-Cox transformations are (A) -0.02, (B) 0.46, (C) 0.18, (D) -0.06, (E) 0.63, (F) 0.75, (G) -0.18, (H) -0.06

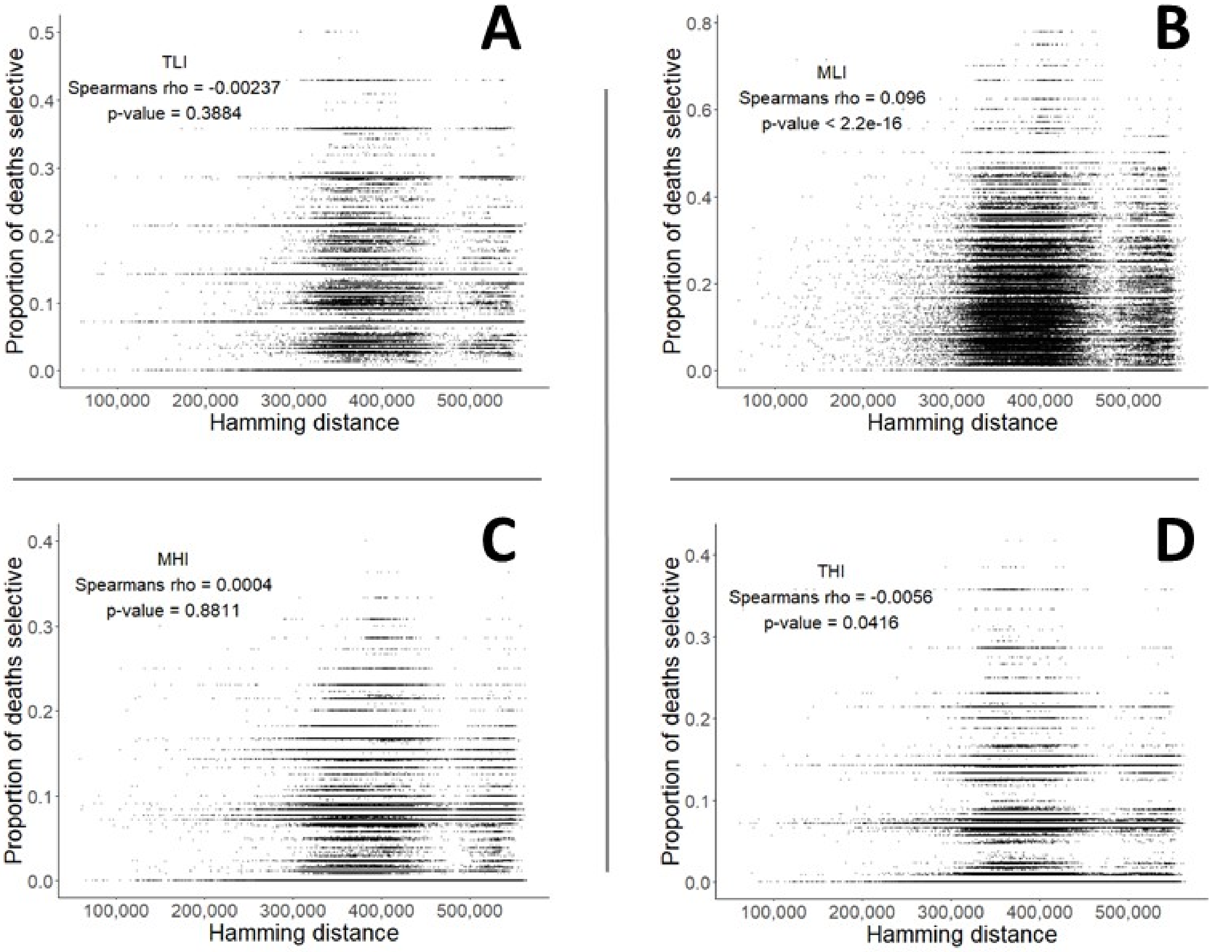


Supplementary Figure 4**.** Hamming distance does not substantially predict the proportion of deaths selective during the life history stage of combined seed and seedling survival at low density.

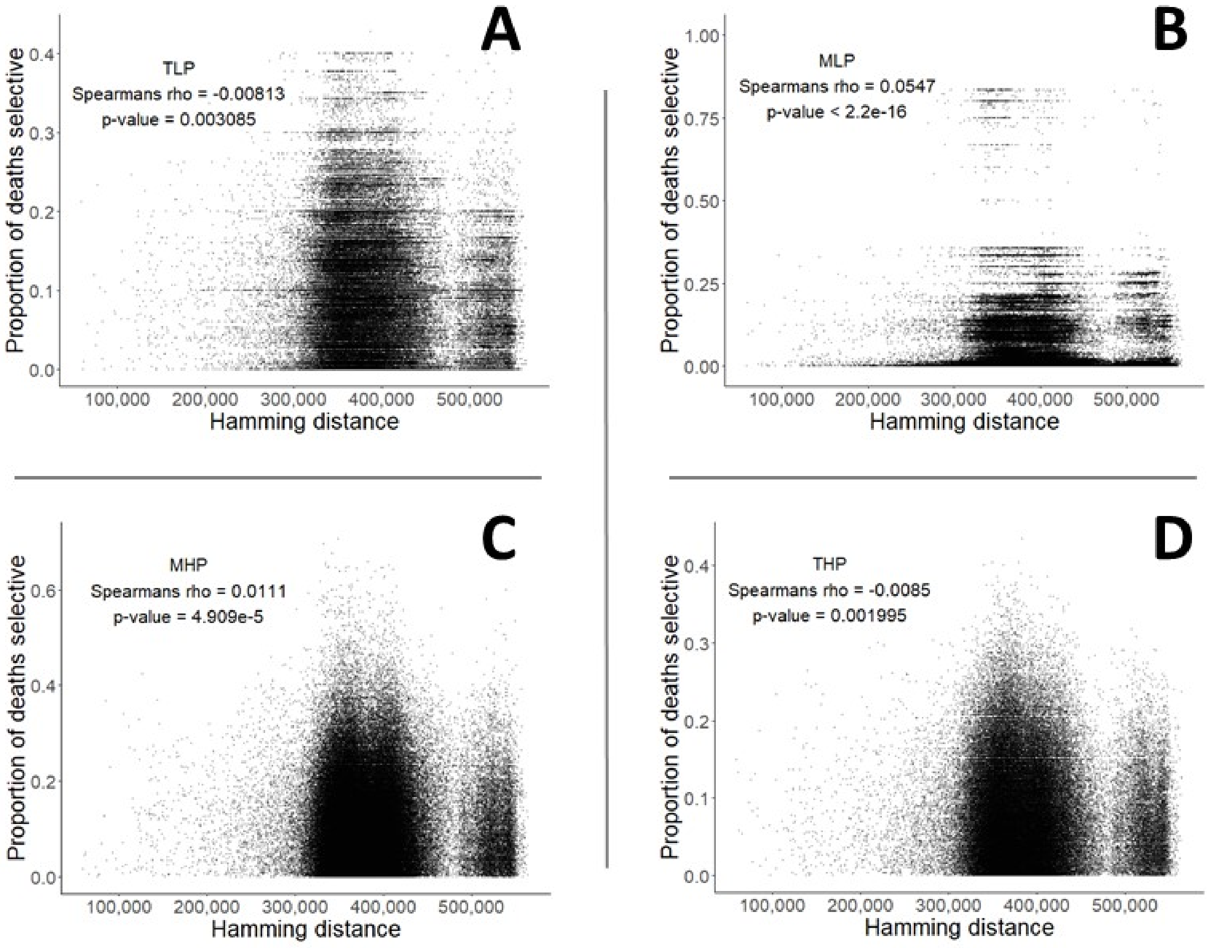


Supplementary Figure 5**.** Hamming distance does not substantially predict the proportion of deaths selective during the life history stage of seedling survival at high density.
